# Fecal pH and redox as functional markers in the premature infant gut microbiome

**DOI:** 10.1101/2023.08.14.553216

**Authors:** Jeffrey Letourneau, LaShawndra Walker, Se Hyang Han, Lawrence A David, Noelle Younge

## Abstract

The infant gut microbiome is a crucial factor in health and development. In preterm infants, altered gut microbiome composition and function have been linked to serious neonatal complications such as necrotizing enterocolitis and sepsis, which can lead to long-term disability. Although many studies have described links between microbiome composition and disease risk, there is a need for biomarkers to identify infants at risk of these complications in practice. In this study, we obtained stool samples from preterm infant participants longitudinally during the first postnatal months, and measured pH and redox, as well as SCFA content and microbiome composition by 16S rRNA gene amplicon sequencing. These outcomes were compared to clinical data to better understand the role of pH and redox in infant gut microbiome development and overall health, and to assess the potential utility of pH and redox as biomarkers. We found that infants born earlier or exposed to antibiotics exhibited increased fecal pH, and that redox potential increased with postnatal age. These differences may be linked to changes in SCFA content, which was correlated with pH and increased with age. Microbiome composition was also related to birth weight, age, pH, and redox. Our findings suggest that pH and redox may serve as biomarkers of metabolic state in the preterm infant gut.

## INTRODUCTION

The gut microbiome plays a crucial role in the health and development of preterm infants. Despite significant improvements in survival of premature infants, alterations to gut microbiome composition and function have been linked to serious neonatal complications such as necrotizing enterocolitis (NEC) and sepsis, which can contribute to long-term disability ^1,2^. Relative to infants born at full term, the preterm infant gut microbiome is characterized by a lack of beneficial commensal bacteria, such as *Bifidobacterium spp*., and an overabundance of hospital-associated pathogens including Enterobacteriaceae^1,3–8^. This dysbiosis of the gut microbiome and its interactions with the immature host intestinal tract and immune system are thought to be central factors in the pathogenesis of these diseases^1,5,6^. While numerous studies have described such links, dysbiosis remains poorly defined and cannot be measured at the bedside in current practice.

Fecal pH and redox potential, which has been less studied, are non-invasive measures of *in situ* conditions in the intestine that may serve as proxies for the microbiome’s metabolic state^19,20^. In adults, pH and redox potential have been shown to be important factors in the maintenance of a healthy gut microbiome. The production of short-chain fatty acids (SCFAs) by gut bacteria results in acidification of the gut, which improves resistance to colonization by pathogens^19,21,22^. However, little is known about the role of pH, redox, and SCFAs in the preterm infant gut microbiome during postnatal colonization. Breastfed infants typically have lower fecal pH due to the predominance of *Bifidobacterium*. Over the past century, the pH in breastfed infants has increased in association with reduction in *Bifidobacterium*^3^. These changes in pH and microbiome overtime are likely multifactorial in nature, related to increased rates of C-section, use of infant formulas, and antibiotic exposure^3^.

This study aims to address this gap in our knowledge and investigate the effects of these factors on the gut microbiome of preterm infants. We hypothesized that serial monitoring of fecal pH and redox potential may provide a means to monitor homeostasis of the gut environment in premature infants. In this study, we obtained stool samples from preterm infant participants (*n* = 11) longitudinally during the first months of life, and measured pH and redox, as well as SCFA content and microbiome composition by 16S rRNA gene amplicon sequencing. These outcomes were compared to demographic and clinical data to better understand the role of pH and redox in infant gut microbiome development and overall health, and to assess the potential utility of pH and redox as biomarkers.

## RESULTS

In order to evaluate the potential for fecal pH and redox potential to serve as biomarkers of metabolic function in the preterm infant gut microbiome, we enrolled 11 infants at the Duke University Hospital Intensive Care Nursery in an observational, longitudinal research study. From fecal samples, pH and redox were measured by benchtop probe, for a total of 77 samples. Additional sample was collected and saved for further analysis of SCFA content and gut microbiome composition. We hypothesized that these variables would be related both to each other and to aspects of infant development such as birth weight and postnatal age.

We first analyzed pH and redox measurements in the context of variables related to birth outcomes, hypothesizing that infants born earlier might have differences in these values. Our results showed that infants with lower birth weight tended to have higher fecal pH (LMM *p* = 0.0046; Fig. 1A), although we found no relationship between birth weight and redox potential (LMM *p* = 0.17; Fig. 1B). Similarly, infants born earlier also had higher pH (LMM *p* = 0.016; Fig. 1C), whereas we did not detect a relationship between birth gestational age and redox (LMM *p* = 0.060; Fig. 1D). The similar results with these two variables are expected, given that infants born earlier tended to weigh less (Spearman correlation *p* = 0.013; Fig. S1A). We hypothesized that redox potential would decline over the first weeks of life given a shift from colonization by aerotolerant to strictly anaerobic bacteria during the first postnatal weeks^6,7,24^. Indeed, while we detected no relationship between postnatal age and pH after controlling for birth weight (LMM *p* = 0.38; Fig. 1E), we did observe that redox potential tended to decrease with age (LMM *p* = 0.0045; Fig. 1F). We also found that pH, but not redox, was related to duration of postnatal antibiotic exposure, even when controlling for birth weight. Specifically, longer antibiotic treatment was associated with higher pH (Fig. 2). These results suggest that developmental stage at birth as well as antibiotic administration may influence fecal pH, and postnatal maturation of the gut microbiome influences redox potential.

**Figure 1.**
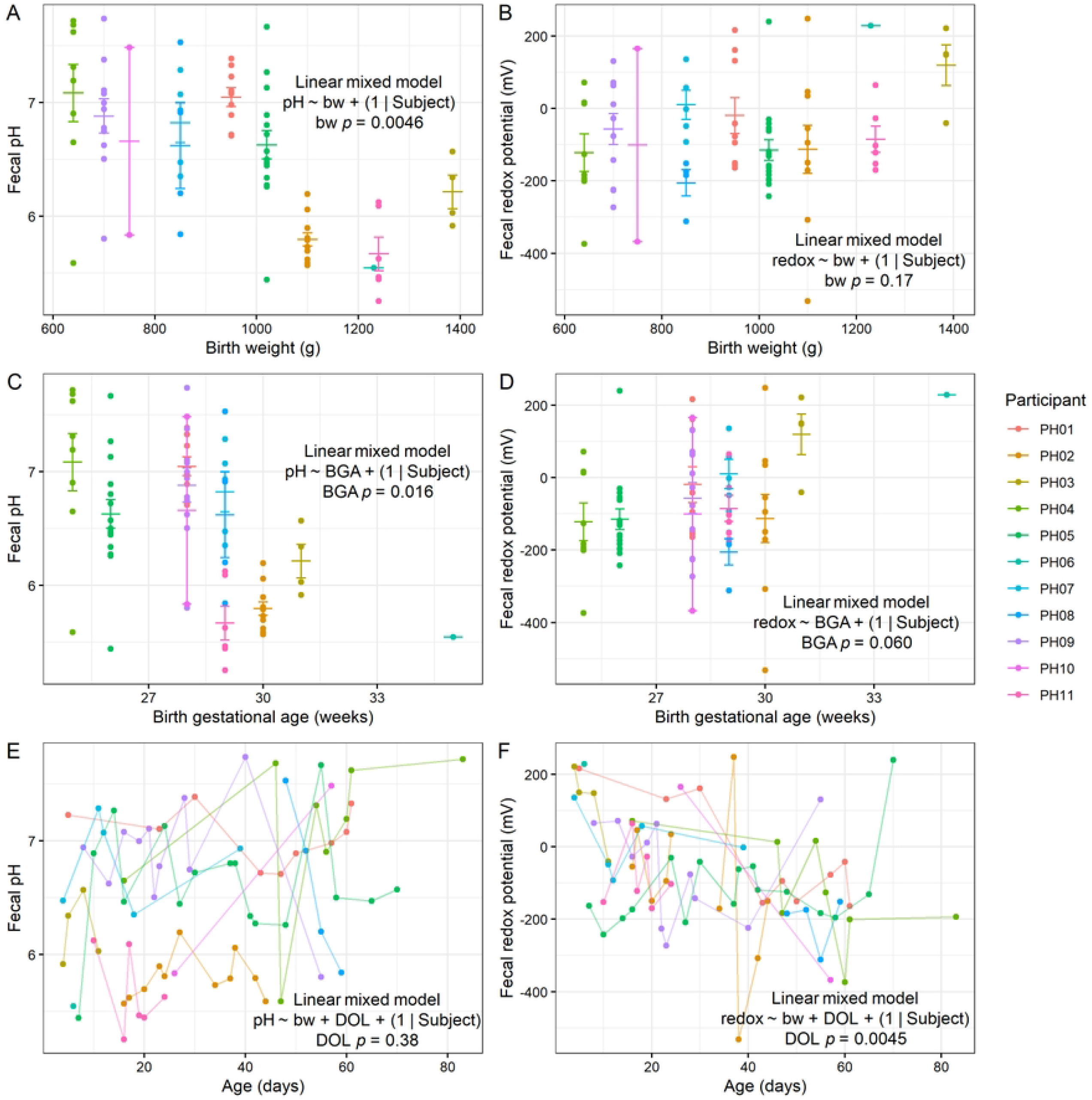
Fecal pH and redox in preterm infants. **A-B**, Relationships between fecal pH (**A**) and redox (**B**) with birth weight. **C-D**, Relationships between fecal pH (**C**) and redox (**D**) with birth gestational age. **E-F**, Relationships between fecal pH (**E**) and redox (**F**) with postnatal age. **A-D**, Mean and standard error shown. **A-F**, LMM results shown (*n* = 11 participants).

**Figure 2.**
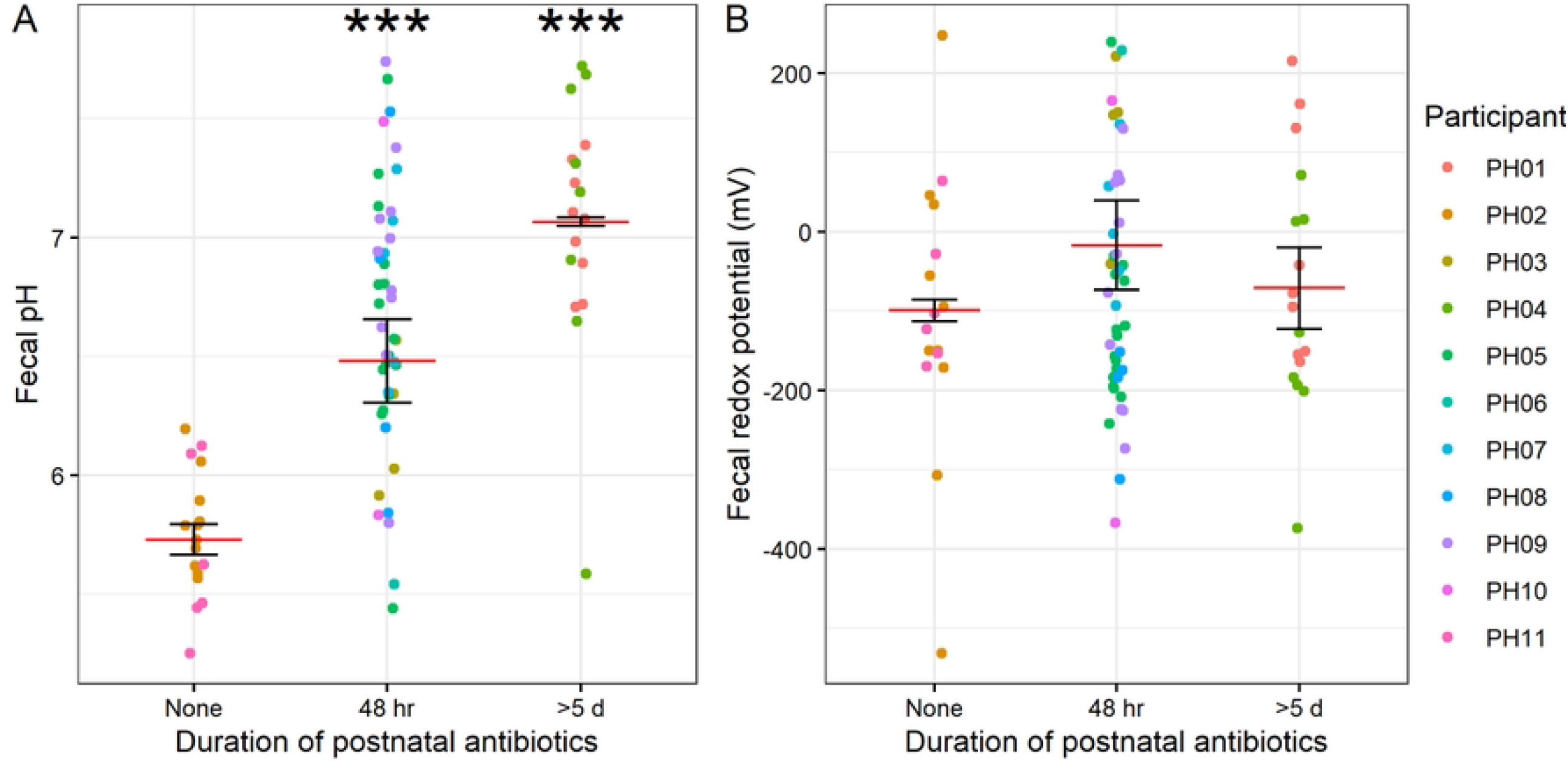
Relationship between antibiotic use and pH and redox. Plots of pH (**A**) and redox (**B**) by duration of postnatal antibiotic treatment. Mean and standard error (of individual participant means) shown. LMM results controlling for birth weight shown (*n* = 11 participants).

To understand what chemical compounds may be driving differences in pH and redox, we measured short-chain fatty acid (SCFA) content by gas chromatography (GC). We detected acetate, butyrate, and propionate, but no detectable levels of isobutyrate, isovalerate, or valerate. As expected, we found that fecal pH was inversely correlated with total SCFA concentration (LMM *p* = 0.0057; Fig. 3A). This relationship was mainly driven by acetate, which was also inversely correlated with pH (LMM *p* = 0.0030; Fig. 3B), although we cannot rule out contributions from other unmeasured compounds such as lactate in determining pH. Furthermore, total SCFA concentration tended to increase with postnatal age (LMM *p* = 0.024; Fig. 3C), and this relationship was mainly driven by acetate (LMM *p* = 0.029; Fig. 3D). Propionate also increased with age (LMM *p* = 0.031; Fig. 3E) and was strongly correlated with current weight (*p* = 0.00047; Fig. 3F). These results suggest that SCFA concentration drives the observed differences in pH in preterm infants, and that concentrations of specific SCFAs increase with age.

**Figure 3.**
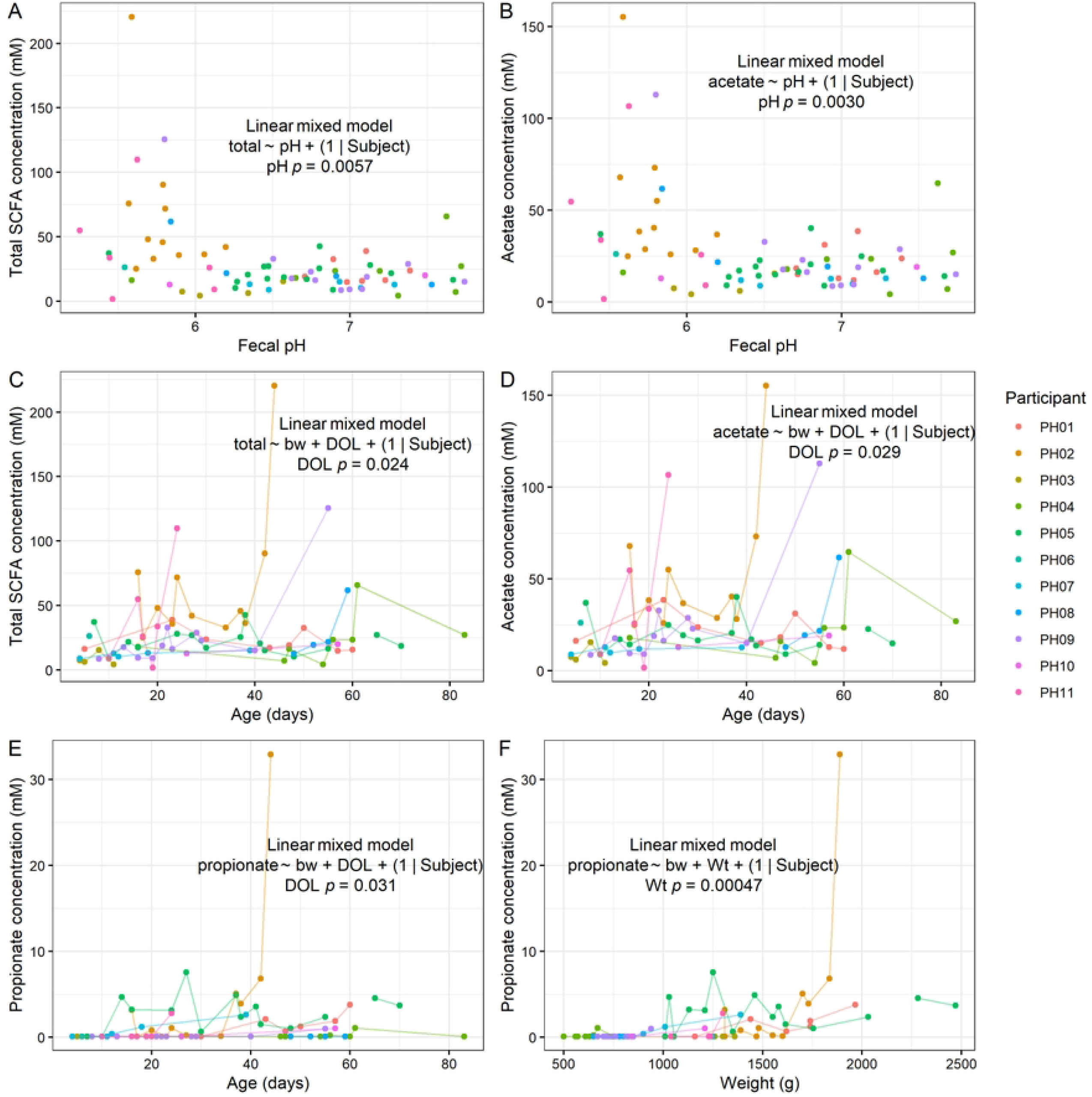
Fecal short-chain fatty acid content in preterm infants. **A-B**, Relationships between total SCFA (**A**) and acetate (**B**) concentrations with fecal pH. **C-D**, Relationships between total SCFA (**C**) and acetate (**D**) concentrations with postnatal age. **E-F**, Relationships between propionate concentration with postnatal age (**E**) and weight (**F**). **A-F**, LMM results shown (*n* = 11 participants).

Finally, we analyzed gut microbiome composition by 16S rRNA gene amplicon sequencing. We found that alpha diversity tended to increase with age, both by observed ASV (amplicon sequence variants) richness (LMM *p* = 0.025) and Shannon index (*p* = 9.1 × 10^-6^; Fig. 4A), although we did not detect a significant relationship between alpha diversity and birth weight (Fig. S2). Microbiome composition also significantly varied by birth weight and day of life (DOL). Interestingly, after controlling for these factors and participant, there was also a significant effect of pH (*p* = 0.0085) and redox (*p* = 0.0054) by PERMANOVA on Bray-Curtis dissimilarity (Fig. 4B). When assessing specific taxonomic differences by ALDEx2 GLM controlling for participant, we identified two ASVs that differed by birth weight (FDR-corrected p < 0.05). These ASVs mapped to *Staphylococcus sp*. (ASV 1), which was negatively correlated with birth weight (ALDEx2 GLM FDR-corrected *p* = 0.0067), while *Escherichia-Shigella sp*. (ASV 4) was positively correlated with birth weight (*p* = 0.033; Fig. 4C). Based on the same model, we also determined that *Staphylococcus sp*. (ASV 1) decreased with postnatal age (*p* = 0.00028; Fig. 4D). These results demonstrate that the microbiome varies in relationship to birth weight and postnatal age, consistent with prior findings.

**Figure 4.**
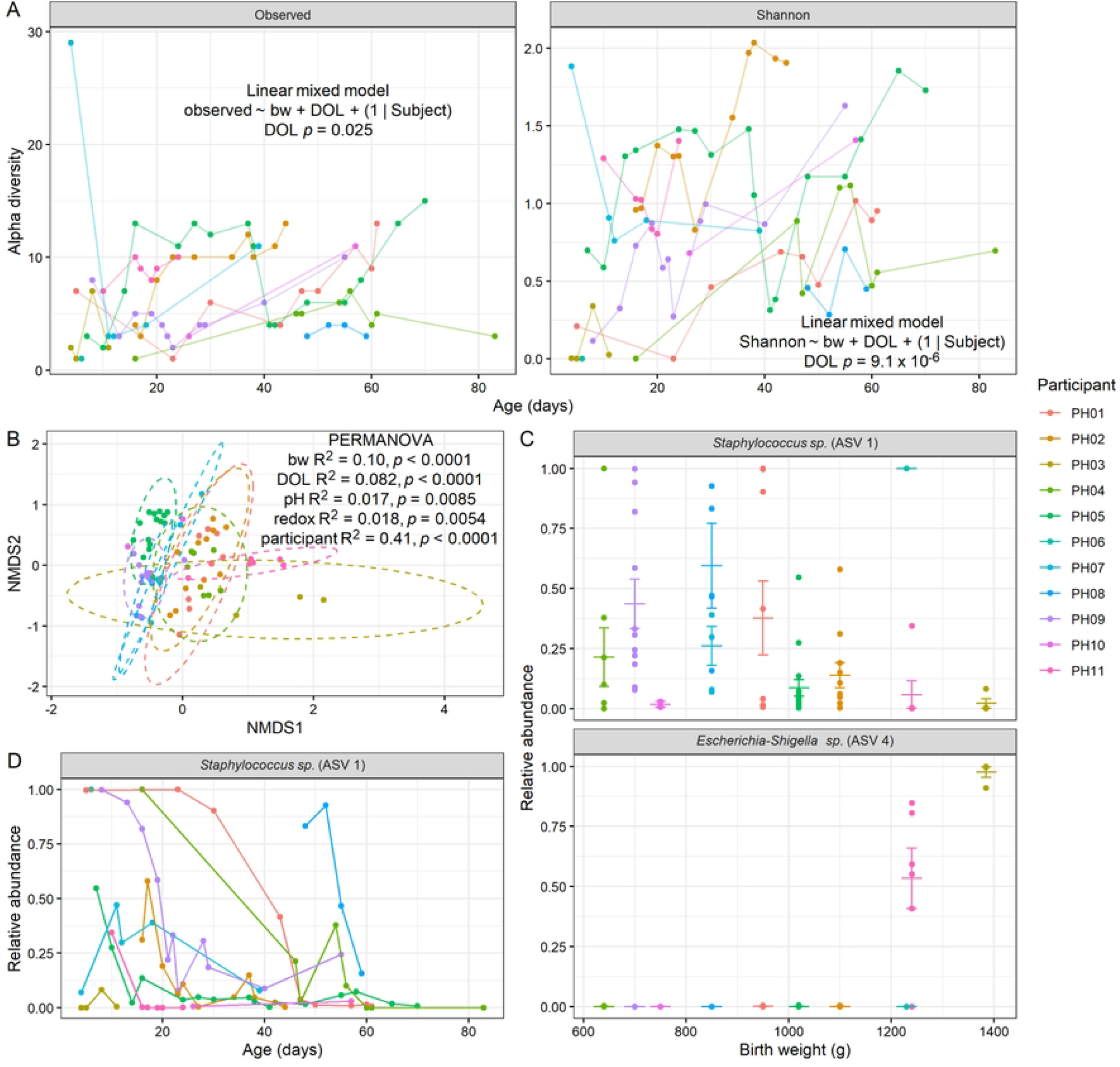
Gut microbiome composition by 16S rRNA gene amplicon sequencing. **A**, Alpha diversity as observed ASV richness and Shannon index, plotted by postnatal age, with results of LMM controlling for birth weight shown. **B**, NMDS ordination plot by Bray-Curtis dissimilarity, with results of PERMANOVA shown. **C**, Two ASVs found by ALDEx2 GLM to significantly differ by birth weight (ASV 1 FDR-corrected *p* = 0.0067, ASV 4 FDR-corrected *p* = 0.033). Mean and standard error shown. **D**, One ASV found to significantly vary by postnatal age (FDR-corrected *p* = 0.00028). **A-D**, (*n* = 11 participants).

## DISCUSSION

The results of this study suggest that pH and redox may serve as biomarkers of metabolic state in the preterm infant gut. Infants born earlier tended to have increased fecal pH, and redox potential increased with postnatal age. These differences may be linked to changes in SCFA content and microbiome composition. Relationships between pH and redox and the adult gut microbiome have been previously described in the literature, but how these dynamics play out in the infant gut is less understood. In this study, we observed that changes in pH and redox were associated with changes in the microbiome composition, although the specific mechanisms underlying these relationships require further investigation.

While numerous studies have characterized the microbiome composition of preterm infants, few have evaluated the relationship of microbiome composition with the biochemical and metabolic environment of the infant gut. However, Our study had several limitations. First, the sample size was small, with only 11 preterm infant participants. On top of this, the resolution of longitudinal sampling varied, and for one participant, we only had a single sample. Nevertheless, this sample size was sufficient to address certain hypotheses relating to birth outcomes and we were sufficiently powered to detect significant relationships between microbial ecological and clinical variables. Follow-up work with a larger cohort size could allow us though to better understand the generalizability of our results and to address further hypotheses. For example, diet (i.e. breast milk vs. different types of formulas) may influence pH and redox, and these readouts of gut metabolic state may in turn influence or reflect risk of diseases such as necrotizing enterocolitis, or conditions such as feeding intolerance and growth failure more broadly. Many studies have determined that maternal breast milk (MBM) is best for extremely premature infants, due to factors such as the presence of human milk oligosaccharides (HMOs) that promote the growth of beneficial bacteria in the infant gut^14–16^. Both diet and antibiotics, which are often necessary to treat infections in preterm infants, can affect microbiome composition and metabolism. Further research is needed to understand how these factors relate to the risk of serious complications such as NEC and sepsis^2,5,18^.

Another related challenge is that of disentangling cause and effect. For example, we found evidence that levels of *Staphylococcus sp*. (ASV 1) were higher in infants with lower birth weight (Fig. 4C), which could perhaps be connected to the increased pH in these lower-weight infants (Fig. 1A). In previous work, we found that antibiotic treatment of mice leads to an abrupt rise in fecal redox potential, with concurrent increases in electron acceptor abundance (e.g., nitrate), oxidative/inflammatory markers (e.g., NOS2 expression), and *Enterobacteriaceae* abundance^23^. Fecal pH has also been reported to decline postnatally as acidic end products are produced by the microbiota during human milk oligosaccharide metabolism. Increases in redox potential and/or fecal pH, may therefore signal a disruption in the environment, such as an increase in inflammation, oxidative stress, or a bloom in facultative anaerobes as is seen in NEC. Follow-up experiments *in vitro* in the absence of host factors may help distinguish cause and effect. Furthermore, a higher sample size would provide the statistical power to assess whether certain microbial taxa relate to pH and/or redox after controlling for other variables.

Despite these limitations, this study provides valuable insights into the role of pH and redox in the development of the gut microbiome in preterm infants, and suggests that these metrics may provide insight into the functional state of the gut microbial ecosystem. Further research is needed to confirm and expand upon these findings, and to explore the potential utility of pH and redox as biomarkers of infant health. Furthermore, understanding how factors such as low birth weight and antibiotic treatment may lead to higher gastrointestinal pH may provide insight into how any negative consequences of this effect may be counteracted. With the increasing recognition of the importance of the gut microbiome in neonatal health and development, such research is critical to improving the care and outcomes of preterm infants through better understanding the underlying microbial ecology.

## MATERIALS AND METHODS

### Participant recruitment and measurement of pH and redox potential

For this prospective single-center pilot study, participants (*n* = 11) were enrolled based on written parental consent, in accordance with the Duke Health Institutional Review Board (IRB) at Duke University under protocol number Pro00100539. A target enrollment of at least 10 infants with repeated sampling was selected to provide sufficient samples to understand the variation in fecal pH and redox over time within and between individuals and to inform subsequent studies of the utility of these measures in this population. Recruitment was conducted (6/1/2019-12/31/2019) by approaching parents (or guardians with capacity to give legally effective consent) of infants in the Duke Intensive Care Nursery (ICN) in person. All infants in the study were born at < 32 weeks gestational age and/or < 1500 g. Infants with congenital gastrointestinal malformations, previous history of necrotizing enterocolitis (NEC), and congenital heart disease were excluded. The median birth gestational age was 28 weeks (range 25-31) and median birth weight was 950 g (range 640-1385). Six infants (55%) were male and 5 (45%) were female. Ten infants (91%) were delivered by C-section, all received human milk feedings, and four (36%) received antenatal antibiotics.

Soiled diapers were collected from enrolled infants until they reached 37 weeks corrected gestational age or were discharged. A benchtop meter (SevenExcellence, Mettler) equipped with pH (InLab Solids Pro ISM) and redox (InLab Redox Micro) sensors were used to take these measurements from the sample. An aliquot of stool was collected and frozen at -80 °C for subsequent analysis.

### SCFA analysis

Due to the small volume of sample available, approximately 100 mg per sample was weighed out into a tube, and a volume of phosphate-buffered saline (PBS) equal to 10 times the sample mass was added, and tubes were vortexed to create a homogenized fecal slurry. SCFAs were then quantified by gas chromatography (GC) as previously described^25^. Briefly, randomized samples were acidified by adding 50 μL 6 N HCl per 1 mL of sample to lower the pH below 3. The mixture was vortexed and then centrifuged at 14,000 × *g* for 5 min at 4 °C. Supernatant was passed through a 0.22 μm spin column filter, and the resulting filtrate was then transferred to a glass autosampler vial (VWR, part 66009-882) equipped with an insert to accommodate the low sample volume. Filtrates were analyzed on an Agilent 7890b gas chromatograph equipped with a flame ionization detector and an Agilent HP-FFAP free fatty-acid column. In this cohort, there were no detectable levels of isobutyrate, valerate, or valerate in any sample. For the remaining SCFAs (acetate, butyrate, and propionate), where the value was below the limit of detection, a pseudocount equal to the lowest value detected for that particular compound was applied for statistical purposes.

### 16S rRNA gene amplicon sequencing

To assess community composition, 16S rRNA gene amplicon sequencing was performed using custom barcoded primers targeting the V4 region of the gene according to previously published protocols^26,27^. The Duke Microbiome Shared Resource (MSR) extracted DNA from stool samples using a Qiagen DNeasy PowerSoil Pro Kit (Qiagen, 47014). Bacterial community composition in isolated DNA samples was characterized by amplification of the V4 variable region of the 16S rRNA gene by polymerase chain reaction using the forward primer 515 and reverse primer 806 following the Earth Microbiome Project protocol (http://www.earthmicrobiome.org/). These primers (515F and 806R) carry unique barcodes that allow for multiplexed sequencing. PCR products concentration was accessed using a Qubit dsDNA HS assay kit (ThermoFisher, Q32854) and a Promega GloMax plate reader. Equimolar 16S rRNA PCR products from all samples were pooled prior to sequencing. Sequencing was performed by the Duke Sequencing and Genomic Technologies shared resource on an Illumina MiSeq instrument configured for 250 base-pair paired-end sequencing runs.

Initial processing of raw sequence data involved custom scripts to create FASTQ files using bcl2fastq v2.20, remove primers using trimmomatic v0.36, and sync barcodes. QIIME2 was used to demultiplex sequenced samples^28^, and DADA2 was used to identify and quantify amplicon sequence variants (ASVs) in our dataset using version 138 of the Silva database^29^. No samples were omitted, as all had over 5000 reads^27^. For analysis of diversity and beta-dissimilarity, all ASVs (95) were included; for analysis of specific taxa by ALDEx2, we retained only those with more than 3 counts in more than 10% of samples, which resulted in 23 ASVs retained.

### Statistics

Statistical analysis was done using custom R scripts, primarily using linear mixed models with participant treated as a mixed effect using the lme4 package, with *p*-values generating using the lmerTest package. For pH, redox, and SCFA, initial analysis of birth statistics (e.g. birth weight) was done by a simple model of the form: pH ∼ birth_weight + (1 | participant). Next, when considering longitudinal variables (e.g. postnatal age), we controlled for birth weight by modeling as: pH ∼ birth_weight + postnatal_age + (1 | participant).

For 16S microbiome composition data, given our findings with the above data, we used a single model of the form: data ∼ birth_weight + postnatal_age + pH + redox + participant. This model formula was used both for PERMANOVA analysis of the Bray-Curtis dissimilarity matrix using the adonis2 function in the vegan package^30^ to assess overall compositional differences, and for a general linear model of the CLR-transformed count data using ALDEx2^31^ to assess differential abundance of specific taxa.

## DATA AND CODE AVAILABILITY

Data for 16S rRNA gene amplicon sequencing will be made publicly accessible via the NCBI SRA prior to publication in the form of demultiplexed reads. Data and code (R scripts) used to generate the figures in this paper and perform all statistical analysis are publicly available on GitHub at the following repository: https://github.com/jrletourneau/infant_microbiome_pH_redox.

## ACKNOWLEDGEMENTS

We would like to thank the Duke Neonatal-Perinatal Research Unit staff who helped with study enrollment and sample collection.

This work was supported by an award from the Duke Microbiome Center, National Institutes of Health grants 1R01DK116187, K23DK129860, the Damon Runyon Cancer Research Foundation, The Hartwell Foundation, and Burroughs Wellcome Fund Investigators in the Pathogenesis of Infectious Disease Award. This study used a high-performance computing facility partially supported by grant 2016-IDG-1013 (HARDAC+: Reproducible HPC for Next-Generation Genomics) from the North Carolina Biotechnology Center.

## AUTHOR CONTRIBUTIONS

Conceptualization: JL, LW, LAD, NY; Data curation: JL, LW, NY; Formal analysis: JL; Funding acquisition: LAD, NY; Investigation: JL, LW, SHH; Software: JL; Visualization: JL; Writing – original draft: JL; Writing – review & editing: JL, LW, SHH, LAD, NY.

